# Climbing fibers selectively recruit disinhibitory interneurons to enhance dendritic calcium signaling in cerebellar Purkinje cells

**DOI:** 10.1101/2025.06.22.660768

**Authors:** Fernando Santos-Valencia, Elizabeth P. Lackey, Aliya Norton, Asem Wardak, Cole S. Gaynor, Sean Ediger, Marie E. Hemelt, Tri M. Nguyen, Wei-Chung Allen Lee, Nicolas Brunel, Court A. Hull, Wade G. Regehr

## Abstract

Climbing fiber (CF) inputs to Purkinje cells (PCs) instruct plasticity and learning in the cerebellum^1–3^. Paradoxically, CFs also excite molecular layer interneurons (MLIs)^4,5^, a cell-type that inhibits PCs and can restrict plasticity and learning^6,7^. However, two types of MLIs with opposing influences have recently been identified: MLI1s inhibit PCs, reduce dendritic calcium signals, and suppress plasticity of granule cell to PC synapses^2,6–9^, whereas MLI2s inhibit MLI1s and disinhibit PCs^8^. To determine how CFs can activate MLIs without also suppressing the PC calcium signals necessary for plasticity and learning, we investigated the specificity of CF inputs onto MLIs. Serial EM reconstructions indicate that CFs contact both MLI subtypes without making conventional synapses, but more CFs contact each MLI2 via more sites with larger contact areas. Slice experiments indicate that CFs preferentially excite MLI2s via glutamate spillover^4,5^. In agreement with these anatomical and slice experiments, *in vivo* Neuropixels recordings show that spontaneous CF activity excites MLI2s, inhibits MLI1s, and disinhibits PCs. In contrast, learning-related sensory stimulation produced more complex responses, driving convergent CF and granule cell inputs that could either activate or suppress MLI1s. This balance was robustly shifted toward MLI1 suppression when CFs were synchronously active, in turn elevating the PC dendritic calcium signals necessary for LTD. These data provide mechanistic insight into why CF synchrony can be highly effective at inducing cerebellar learning^2,3^ by revealing a critical disinhibitory circuit that allows CFs to act through MLIs to enhance PC dendritic calcium signals necessary for plasticity.

---

CFs make powerful excitatory synapses onto PCs that elevate dendritic calcium and can induce long-term synaptic depression (LTD) at granule cell (GrC) to PC synapses^9–12^. CFs also regulate the output of the cerebellar cortex by generating complex spikes (PC_CS_), that are followed by a pause in the normally high simple spike firing of PCs (PC_SS_)^13,14^. In addition to directly influencing PCs, CFs excite MLIs that can inhibit PCs to regulate the output of the cerebellar cortex. This inhibition can gate GrC-PC plasticity by reducing calcium signals in PC dendrites and preventing LTD, a form of plasticity thought to be necessary for many forms of cerebellar learning^6,9^. This presents an intriguing dilemma: how can CFs facilitate learning when they also activate interneurons that suppress the dendritic calcium signals in PCs necessary for LTD?

The recent discovery of two MLI subtypes^15^ has the potential to resolve this conundrum. Previous electrophysiological studies of CF-MLI synapses did not discriminate between MLI subtypes^4,5,16–19^, and it is not known if CFs influence MLI1s and MLI2s in the same way. This has important functional consequences because MLI1s are electrically coupled to each other, fire synchronously, and preferentially inhibit PCs, whereas MLI2s are not electrically coupled, preferentially inhibit MLI1s and disinhibit PCs^8^. This suggests that MLI1s can suppress LTD and MLI2s can promote LTD at GrC to PC synapses, together gating CF-induced synaptic plasticity and learning^20^. However, it is not clear how this gating could operate at the circuit level, as there is no known mechanism for differentially recruiting MLI subtypes. We therefore tested how CFs engage MLI subtypes, and how CF activation of MLIs influences PC spiking and dendritic calcium signaling.

## CFs preferentially contact MLI2s via unconventional synapses

To provide insight into how CFs could activate MLIs, we performed serial EM reconstructions of CFs (*blue*), PCs (*grey*), MLI1s (*purple*) and MLI2s (*green*) identified as described previously^8^ (**Fig 1a**). CF boutons that contact either type of MLI are large and contain many vesicles (**Fig. 1a**, *insets*). We find that each CF contacts about the same number of MLI1s and MLI2s (4.80 ± 0.68 MLI1s and 4.90 ± 0.89 MLI2s; **Fig. 1b-c**, *bottom*). However, each CF makes approximately 2.4 (16.6/6.8) times as many contacts onto MLI2s (**Fig. 1b-c**, *top*). This difference is even more striking given that there are 3-3.5 times as many MLI1s as MLI2s^8,15^. From the perspective of individual MLIs, MLI1s were contacted by 0.40 ± 0.16 CFs that made a total of 0.50 ± 0.22 contacts (**Fig 1d**, *bottom*), and MLI2s were contacted by 1.80 ± 0.25 CFs that made a total of 4.50 ± 0.85 contacts (**Fig 1d**, *top*, and **Fig 1e**).

**Fig. 1.**
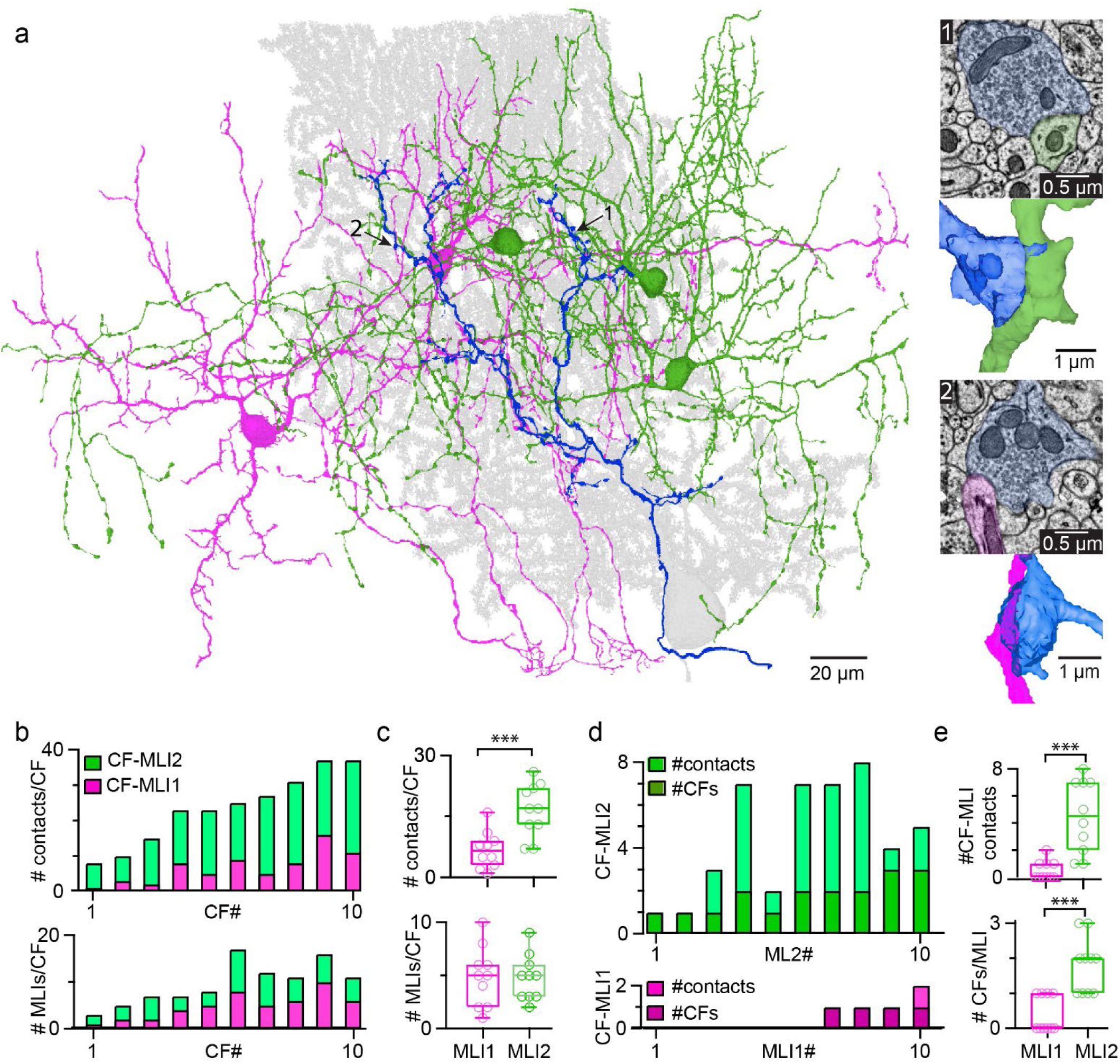
EM reconstructions of CF to MLI1 and MLI2 synapses. **a.** Image of a reconstructed CF (*blue*), the corresponding PC (*grey*) and an MLI1 (*purple*) and MLI2 (*green*) that are contacted by the CF. Arrows indicate two points of contact. inset 1: (*top*) an EM section showing the point of contact between a CF and an MLI2. inset 1: (*bottom*) a reconstruction of the point of contact between a CF and an MLI2. inset 2: same as inset 1 but for an MLI1. **b.** Summary of contacts made by 10 CFs onto MLIs, with the number of MLIs contacted (*bottom*) and the total number of contacts made by the CF (*top*) summarized for both MLI subtypes. **c.** Summary of the numbers of MLIs and contact sites for each CF. **d.** Summary of the number of CFs and CF contact sites for 10 MLI1s (*purple*) and 10 MLI2s (*green*). **e.** Summary of the number of CFs and CF contact sites for MLI subtypes.

Consistent with previous anatomical and physiological work suggesting that CF to MLI input is mediated by unconventional synapses and spillover^4,5,19,21^ (although see^22^), we find that CF boutons onto both types of MLIs are not associated with a cluster of docked vesicles near a postsynaptic density on the opposing membrane (**Fig. 2ab, Extended Data Fig. 1ab**). This contrasts with GrC to MLI2 synapses, as well as CF to PC synapses, that have a cluster of docked vesicles associated with a postsynaptic density (PSD) that is characteristic of a fast, conventional synapse (**Fig. 2ab**). Motivated by the observation that CF stimulation activates AMPARs associated with nearby GrC to MLI synapses^19^, we reconstructed GrC parallel fiber axons and their associated PSDs on a dendrite near CF-MLI2 contacts (**Fig. 2cd, Extended Data Fig. 1c**) and near CF-MLI1 contacts (**Fig. 2e-f, Extended Data Fig. 1d**). These reconstructions show substantial differences in CF contacts onto MLI1s and MLI2s. CF-MLI2 contact sites are larger than CF-MLI1 contact sites (**Fig. 2g**, 1.60 ± 0.74 vs 0.66 ± 0.33 μm^2^), more GrC PSDs are within 1 μm of each CF-MLI2 contact than CF-MLI1 contacts (**Fig. 2h**, 2.9 ± 1.3 vs 1.20 ± 0.78), and the close proximity of CF-MLI2 contacts to so many GrC PSDs is expected to facilitates spillover responses^19,23,24^. Thus, CFs are predicted to more strongly activate MLI2s than MLI1s, because MLI2s are contacted by a greater number of CFs, each CF makes more contacts onto each MLI2, and individual contacts onto MLI2s are larger and in closer proximity to more GrC PSDs.

**Fig. 2.**
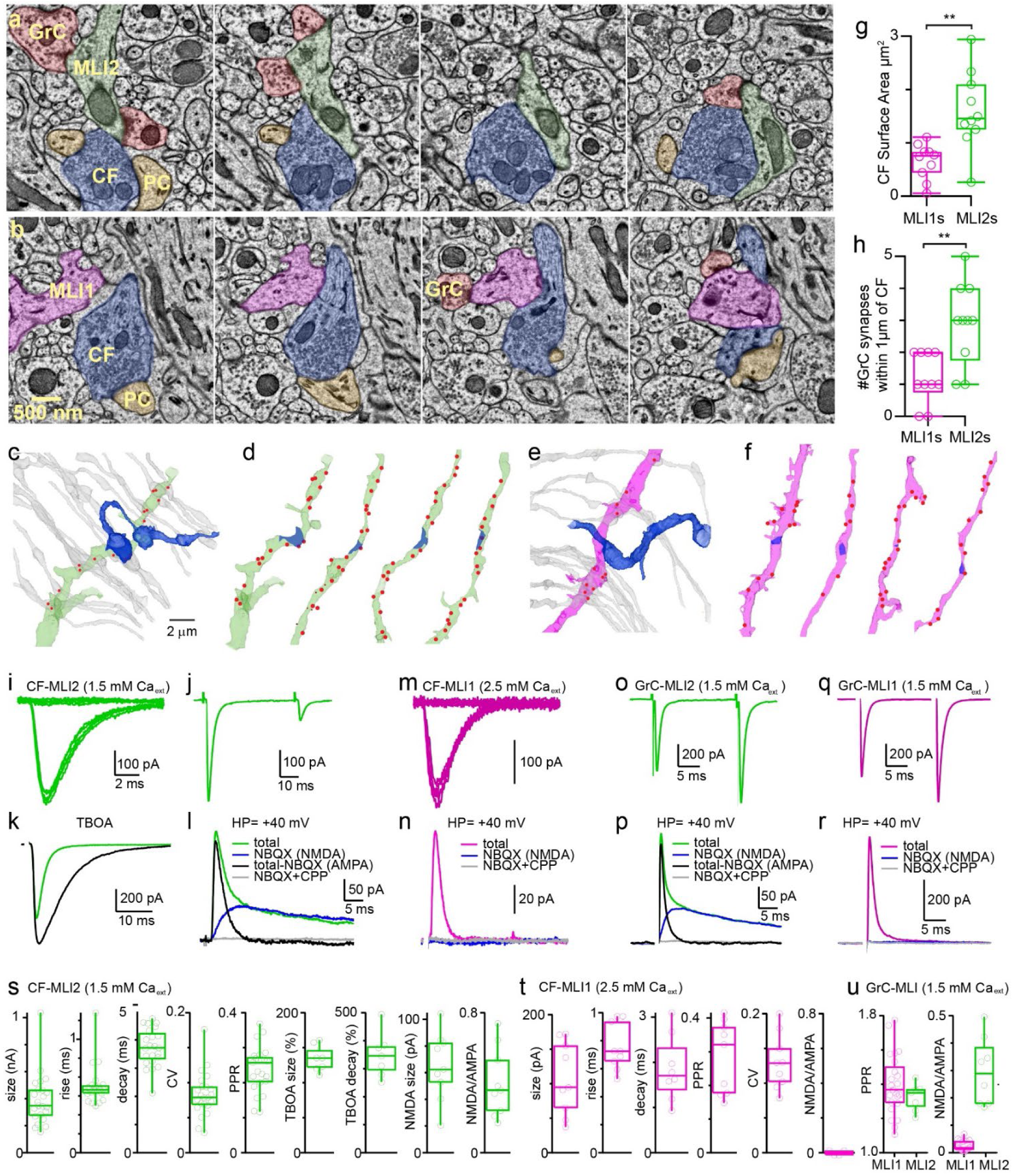
CF-MLI2 responses are strong and mediated by glutamate spillover. **a.** A series of EM sections shows a CF (*blue*) contacting an MLI2 (*green*). Docked vesicles across from PSDs were not observed at CF-MLI2 contacts, but were apparent at GrC (*red*) to MLI2 synapses and CF to PC (*orange*) synapses. **b.** A series of EM sections shows a CF (*blue*) contacting an MLI1 (*purple*). Docked vesicles across from PSDs were not observed at CF-MLI1 contacts, but were apparent at GrC (*red*) to MLI1 synapses and CF to PC (*orange*) synapses. **c.** *left* An EM reconstruction of a CF (*blue*) to MLI2 (*green*) contact, granule cell parallel fibers (grey), and granule cell-MLI2 PSDs (*red*). **d.** Four additional MLI2 dendrites are shown as in **c**, with CF-MLI2 contacts (*blue*), and granule cell-MLI2 PSDs (*red*). **e-f**) As in **c-d**, but for MLI1s (*purple*). **g.** Summary of CF-MLI contact areas. **h.** Summary of the number of GrC PSDs within one micrometer of the center of a CF-MLI contact site. **i-l.** CF-MLI2 EPSCs (1.5 mM Ca_ext_) **i.** CF-MLI2 EPSCs were all-or-none as illustrated by 10 consecutive responses evoked by the same intensity stimulation. **j.** CF-MLI2 EPSC paired-pulse depression. **k.** The effect of blocking glutamate uptake (50 μM TBOA, *black*) on a CF-MLI2 EPSC. **l.** A CF-MLI2 EPSC was recorded at +40 mV (*green*), followed by successive application of the AMPAR antagonist NBQX (5 μM, *blue*), which was then coapplied with the NMDAR antagonist CPP (4 μM, *grey*). The AMPA component (*black*) was determined by subtracting the trace in the presence of NBQX from the initial trace. **m, n.** CF-MLI1 EPSCs (2.5 mM Ca_ext_) **m.** CF-MLI1 EPSCs were all-or-none as illustrated by 10 consecutive responses evoked by the same intensity stimulation. **n.** As in **l** but for GrC-MLI2 EPSCs. **o, p.** GrC-MLI2 EPSCs (1.5 mM Ca_ext_). **o.** Paired-pulse facilitation of GrC-MLI2 synapses. **p.** As in l but for GrC-MLI2 synapses. **q, r.** As in **o, p,** but for GrC-MLI1 EPSCs. **s-u.** Summary of CF and GrC synaptic properties for MLI2s and MLI1s.

## CFs robustly excite MLI2s via glutamate spillover

To determine the impact of CF synapses onto MLI subtypes, we made whole-cell recordings from MLI1s and MLI2s in acute brain slices. In physiological external calcium (1.5 mM Ca_ext_), CF-MLI2 synapses were observed in 25 of 26 MLI2s, and their properties are summarized in **Fig. 2s**. CF-MLI2 synapses are large, slow, all-or-none (**Fig. 2i**), had pronounced paired-pulse depression (**Fig. 2j**), and are slowed by blocking glutamate uptake with TBOA (**Fig. 2k**, n=8). These properties are consistent with CF spillover synapses in which glutamate diffuses to distant locations^4^. Previous studies showed that some CF-MLI synapses can have prominent NMDA components^4,5^, and we find that the NMDA component is large and reliably observed at CF-MLI2 synapses (**Fig. 2l, 2s**, n=9). In contrast, CF-MLI1 synapses were difficult to observe in 1.5 mM Ca_ext_ and were well isolated in 5% (1/20) MLI1s. However, when synaptic strength was increased by elevating Ca_ext_ to 2.5 mM, CF-MLI1 EPSCs were observed in 61% (25/41) of MLI1s and well-isolated in 27% of MLI1s. CF-MLI1 synapses are also all-or-none, depressed, and have a slow decay time (**Fig. 2m, t**). Surprisingly, CF-MLI1 synapses are mediated exclusively by AMPARs, and an NMDA component is not apparent (**Fig. 2n, 2t**, n=4). These results indicate that CF-MLI2 synapses are much stronger than CF-MLI1 synapses, and CF-MLI2 synapses are mediated by both NMDARs and AMPARs, whereas CF-MLI1 synapses are mediated almost exclusively by AMPARs.

If CF responses in MLI2s are mediated by glutamate spillover activation of receptors at GrC-MLI2 synapses^19^, then a prominent NMDA component should be present at GrC-MLI2 synapses but not at GrC-MLI1 synapses. Previous studies have shown that an NMDA component is present at GrC to MLI synapses, but in some cases the component is very small and in others it is prominent^25–27^. We find that both GrC-MLI1 (n=23) and GrC-MLI2 synapses (n=8) facilitated (**Fig. 2m, q, u**), but a prominent NMDA component is only observed at GrC-MLI2 synapses (**Fig. 2n, r, u**). These results are consistent with snRNAseq studies^15^ and *in situs* (**Extended Data Fig. 2**) in which *Grin2b* (the Nr2b subunit) is expressed at much higher levels in MLI2s (*Nxph1^+^*) than in MLI1s (*Sorcs3^+^*). When combined with the finding that NMDARs in MLIs are essential to LTD and motor learning^28^, the preferential contribution of NMDARs to MLI2 responses suggests that NMDARs in MLI2s are likely crucial for motor learning.

## Spontaneous activation of CFs regulates firing of MLIs and disinhibition of PCs

We used serial EM to further explore the influence of CF activation on populations of MLIs in the molecular layer. Our working hypothesis is that CF contacts onto MLIs bring CFs into close proximity to GrC to MLI synapses to mediate CF-MLI EPSCs via spillover activation of PSDs associated with GrCs^19^. This suggests that even though CF-MLI contacts are not conventional fast synapses, the contact sites provide insight into how activity may be influenced by CFs. In addition to considering direct CF-MLI contacts (**Fig. 3a**), it is important to consider the many MLI1s that are inhibited by the MLI2s that are contacted by the CF (**Fig. 3b**). In the example of **Fig. 3ab**, the CF contacts 5 MLI1s and 5 MLI2s (**Fig. 3a**), and these MLI2s in turn inhibit 33 MLI1s (**Fig. 3b**). MLI1s that are predicted to be disynaptically inhibited (CF-MLI2-MLI1) in turn synapse onto many PCs. We therefore reconstructed the circuitry to predict the effects of CF activation on PCs. For an example PC directly contacted by the CF, we find that 149 CF-MLI1-PC synapses provide disynaptic inhibition, and 543 CF-MLI2-MLI1-PC synapses provide disinhibition (**Fig. 3c**, *left*). For a neighboring PC, we find that 95 CF-MLI1-PC synapses provide disynaptic inhibition, and 517 CF-MLI2-MLI1-PC synapses provide disinhibition (**Fig. 3c**, *right*). Thus, our reconstructions suggest CF-MLI activation will lead to stronger disinhibition than inhibition for both directly connected and neighboring PCs (**Fig. 3d, Extended Data Fig. 3**).

**Fig. 3.**
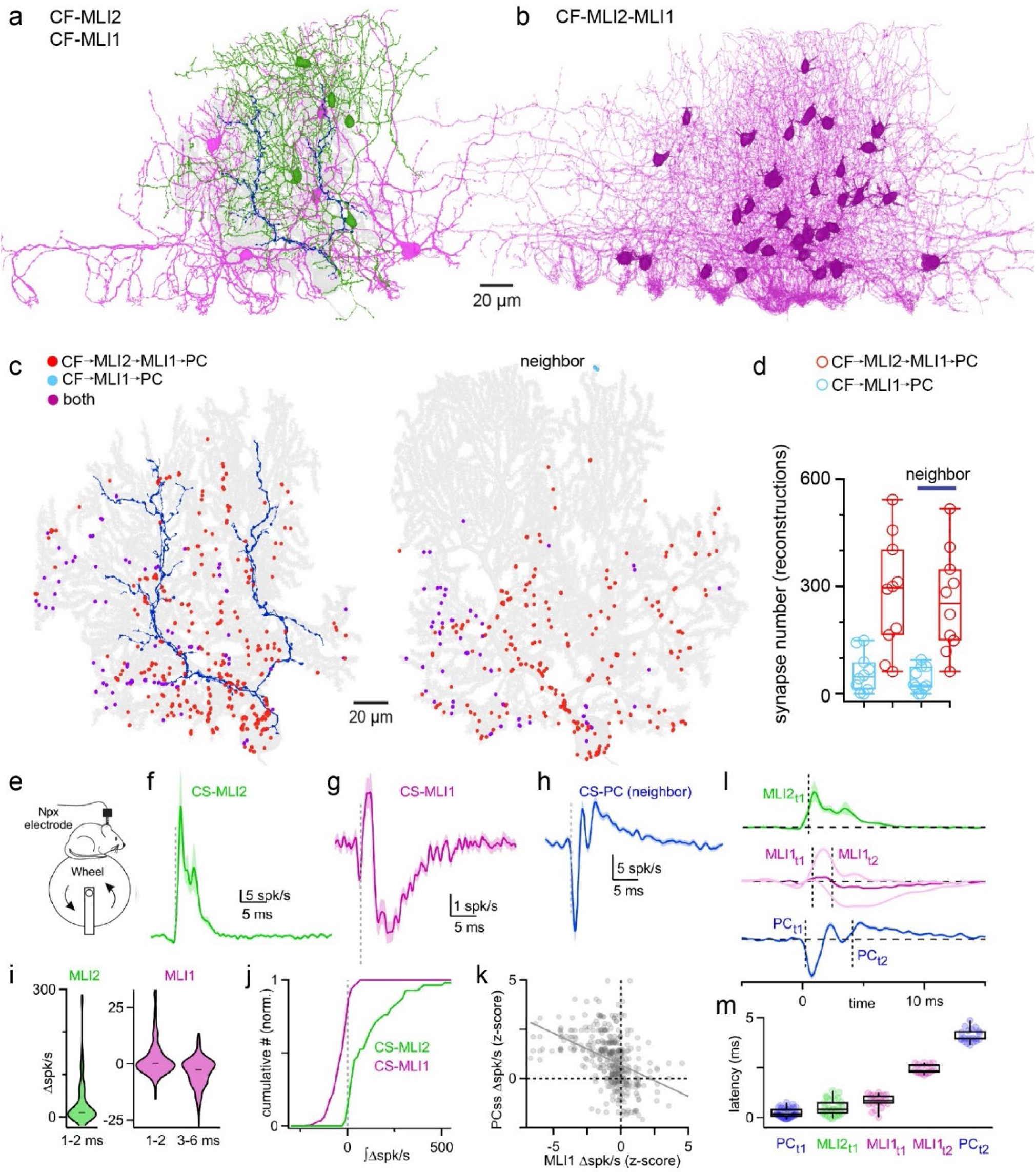
CF activation of MLI circuitry. **a.** A CF (*blue*) synapsing onto a PC (*grey*) is shown along with the MLI1s (*purple*) and MLI2s (*green*) contacted by the CF. **b.** CFs activate MLI2s that inhibit MLI1s that in turn inhibit the PC (PC1) that is contacted by the CF. MLI1s that are part of this circuit are shown. **c.** EM reconstructions were used to evaluate the circuitry influenced by CF activation. (*left*) For a PC directly excited by a CF, inhibitory synapses made by MLI1s that are contacted by the CF (CF-MLI1-PC1, light blue), MLI1-PC1 synapses that are CF-MLI2-MLI1-PC1 circuit (*red*), and MLI1-PC synapses for MLIs that are both directly contacted by CFs, and part of the CF-MLI2-MLI1-PC1 circuit (*purple*). (*right*) As above, but for a nearest neighbor PC. **d.** Summary of predicted CF-MLI1 and CF-MLI2-MLI1 inputs onto PCs and neighboring PCs as in **c**. **e.** Schematic showing the experimental setup for the *in vivo* recordings in (**f-m**). **f.** Average CS-MLI2 CCGs for spontaneous CSs. **g.** Average CS-MLI1 CCGs for spontaneous CSs. **h.** Average CS-PC CCG for neighboring PCs for spontaneous CSs. **i.** Summary of the peak increases in firing for CS-MLI2s and CS-MLI1 CCGs, and the maximal decrease in firing for CS-MLI1 CCGs. **j.** The effects of CSs on the firing of MLIs is summarized in normalized plots of the cumulative integral of the changes in the number of spikes for all CS-MLI CCGs. **k.** The relationship between the peak z-scores for the changes in PC_SS_ firing rates and MLI1 firings rates is shown for all nearby MLI1-PC pairs (R^2^=0.19). **l.** Plots of average CS-MLI2, CS-MLI1, and CS-PC_SS_ CCGs as in **f**-**h** showing the timing of various components. The dark purple line is the average of all CS-MLI1 CCGs, and the two faint purple lines are averages of subsets of CS-MLI1 CCGs with either a Z-score > 10 (1-2 ms), or < −10 (3-6 ms). Vertical lines indicate average latencies in **m**. **m.** Summary of the latencies for CS-MLI2s, CS-MLI1s increases in firing, CS-MLI1s decreases in firing, CS-PC_SS_ ephaptic suppression, CS-PC_SS_ disinhibition.

Reconstructions and slice recordings predict that CF activation will lead to a large increase in the firing of MLI2s, a small increase followed by a large decrease in MLI1 firing, and disinhibition of PC firing. We tested these predictions using high density silicon probe recordings (Neuropixels) in behaving mice (**Fig. 3e-m**). Spontaneous CF activation reliably drives complex spikes in PCs (PC_CS_), which serve as a proxy for CF activity. PC_CS_-MLI2 cross-correlograms (CCGs) reveals a large increase in MLI2 firing just after PC_CS_ onset (**Fig. 3f,i**), whereas PC_CS_-MLI1 CCGs show only a small increase in MLI1 firing followed by a decrease in firing after PC_CS_ onset (**Fig. 3g,i**). For 77% (43/56) of the PC_CS_-MLI2 CCGs there is a short latency increase in firing (Z-score > 4 for 1-2 ms after the PC_CS_), and for PC_CS_-MLI1 CCGs 25% (47/190) were accompanied by a short-latency increase in firing (Z-score > 4 for 1-2 ms after the PC_CS_) and 39% (74/190) by a longer-latency decrease in firing (Z-score < −4 for 3-6 ms after the PC_CS_). The pronounced differences in the PC_CS_-MLI1 and PC_CS_-MLI2 CCGs is illustrated by quantifying the cumulative change in spikes associated with the PC_CS_ for each PC_CS_-MLI pair: a PC_CS_ is associated with an increase in spiking (Z-score > 4) in 30% of MLI1s and 93% of MLI2s, and a decrease in spiking (Z-score < −4) in 57% of MLI1s and 2% of MLI2s (**Fig. 3j**). It is difficult to assess the effect of CFs activation of MLIs on the PC targeted by the CF because a PC_CS_ is followed by a pause in simple spiking (PC_SS_) produced at least in part by intrinsic conductances^29^. We therefore instead evaluate the interaction between PC_CS_ and PC_SS_ in neighboring PCs and find rapid decreases in spiking followed by slower elevations in spiking (**Fig. 3h**).

The timing of the effects of PC_CS_ on the firing of nearby cells is shown by aligned average CCGs for different cell types with dashed vertical lines showing the components we evaluated (**Fig. 3l**), and by a summary of the timing of the components for individual CCGs with large Z-scores (**Fig. 3m**). PC_CS_ -MLI2 CCGs show that the latency of MLI2 activation from PC_CS_ onset is ∼ 0.50 ms. The average of all PC_CS_ -MLI1 CCGs shows an initial increase in firing followed by a decrease in firing (**Fig. 3g**, **Fig. 3l**, *dark purple*). A subset of PC_CS_ -MLI1 CCGs showed prominent short latency increases in firing (∼0.83 ms) and another showed longer latency suppression (∼ 2.4 ms) (**Fig. 3l**, *light purple traces*). For PC_CS_ -PC_SS_ CCGs there is a very rapid decrease in firing (0.23 ms latency) followed by a much longer latency (4.1 ms) increase in firing (**Fig. 3l**, *blue*). The rapid decrease of PC_SS_ rate after a PC_CS_ is the result of ephaptic coupling in which the CFs produces an extracellular signal that rapidly inhibits neighboring PCs^30^ (**Fig. 3m**); PC_CS_ then rapidly excite MLI2s and a small fraction of MLI1s, ∼ 1.9 ms later MLI2s inhibit MLIs, and after an additional ∼1.7 ms there is an increase in PC_SS_ firing that is consistent with disinhibition mediated by the CF-MLI2-MLI1-PC pathway. If the increase in PC_SS_ firing is a consequence of disinhibition involving MLI1s, then PC_SS_ spiking should be inversely correlated with the suppression of nearby MLI1s. We tested this possibility by plotting z-scores of the firing rate of connected MLI1s to PC_SS_ to in response to the same PC_CS_ and find that, as predicted, large decreases in MLI1 firing are correlated with large increases in PC_SS_ firing (**Fig. 3k**).

## Sensory input has heterogeneous effects on MLI1s

We have shown that spontaneous CF inputs preferentially drive MLI2s to inhibit MIL1s and disinhibit PCs, but cerebellar learning relies on sensory-driven CF inputs, which can differ from spontaneous CF inputs^31,32^. To determine how MLIs are recruited by learning-related inputs, we delivered a periocular airpuff stimulus of the kind typically used as the unconditioned stimulus for cerebellar-dependent eyelid conditioning, and thought to be carried to the cerebellar cortex by the CF pathway as the instructive signal. We then evaluated how this sensory stimulus influences the firing of circuit elements in the cerebellar cortex. Experiments were performed in locomoting mice because sensory-driven PC_CS_ are more effective at inducing learning during locomotion than when mice are stationary^33^. As expected, airpuffs reliably increase PC_CS_ firing (50/244) (**Fig.4c**). Surprisingly, while airpuffs also strongly excite a subset of MLI2s (6/7, **Fig. 4d**), they had only a minor influence on average MLI1 firing (**Fig. 4e**) and PC_SS_ firing (**Fig. 4b**). To determine why MLI2s may be less effective at suppressing MLI1s during sensory-evoked activity, we measured airpuff-evoked responses arising from the mossy fibers (MF), the input pathway classically thought to carry the conditioning stimulus information. We find that airpuffs also elevate MF firing (44/260, **Fig. 4a**), suggesting that this secondary input may be competing with the disinhibitory effects of the CF-MLI2-MLI1 pathway.

**Fig. 4.**
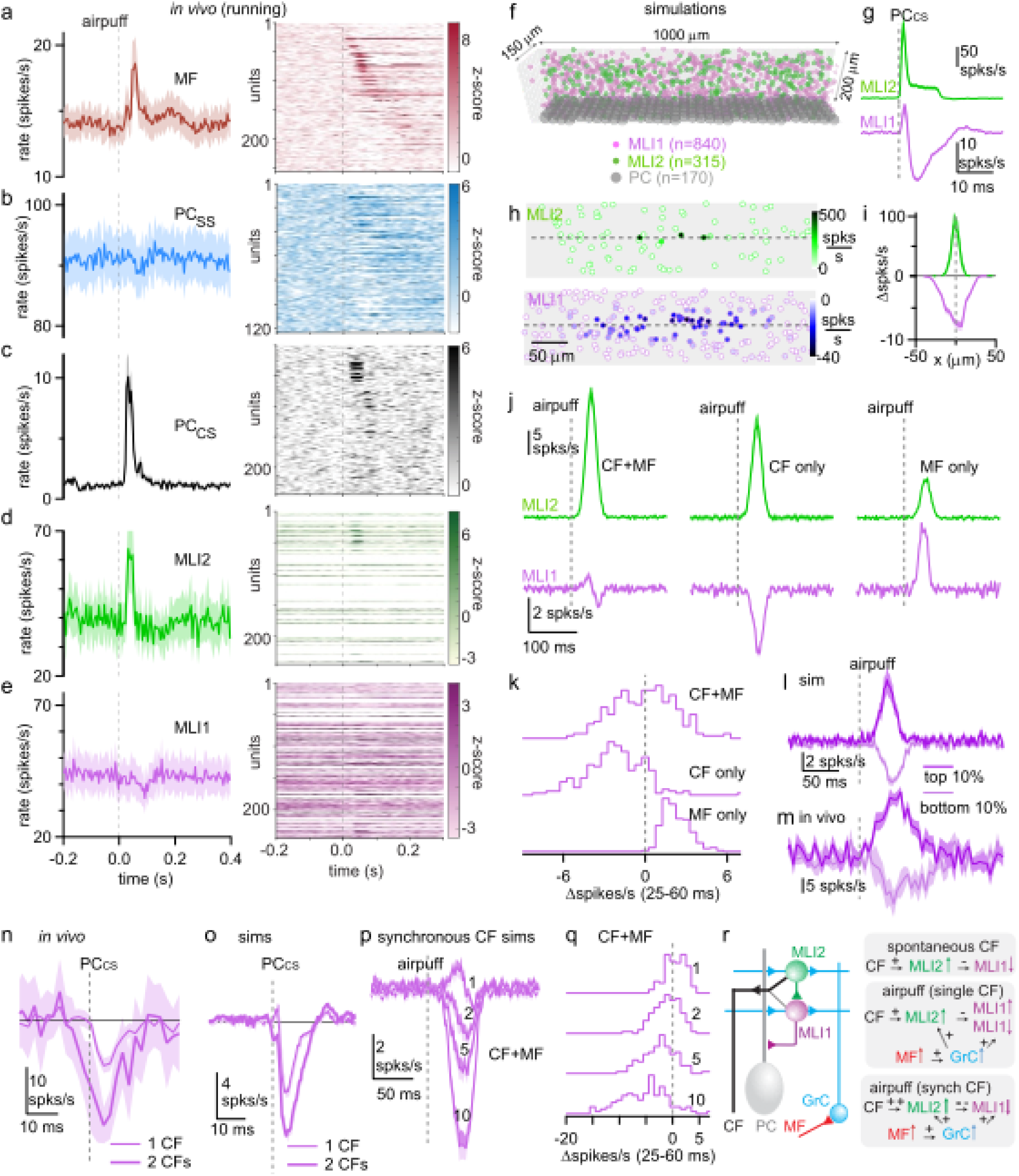
Sensory-driven MLI responses and the effects of CF synchrony. **a.** Airpuff-evoked responses of mossy fibers (256 units) during running. Averaged PSTH of all mossy fibers (left) and z-scored heatmap (right) sorted by latency to peak response. **b-c.** Same as a but for PCss of neighboring PCs (125 units) and for PCs with an airpuff activated CS PCcs (50 of 244 units). **d.** Airpuff responses of MLI2s (12 of 18 units) connected to an airpuff driven PCcs during stationary periods. Averaged PSTH (left) and z-scored heatmap of the averaged response of MLI2s connected to a given PCcs in **c**. This procedure keeps the same sorting as the heatmap in **c**., allowing MLIs to be represented on multiple rows if they are connected to more than one CF. In cases in which no PCcs-MLI2 pair exists, a white line was added to the heatmap. **e.** Same as **d** but for MLI1s (50 of 112 units). **f.** Schematic of model used for simulations in (**f-l**). The model is constrained by EM reconstructions with MLI properties and synaptic connections constrained by a combination of em reconstructions, slice recordings and *in vivo* recordings. **g.** Average responses of MLI1s and MLI2s evoked by spontaneous CF activation. **h.** Spatial maps of individual MLI1s and MLI2s evoked by spontaneous firing of a CF. All MLIs are sown in these top-down views and symbol fills indicate their changes in firing rates. The plane containing the CF is approximated by the dashed line. **i.** Average CF-evoked MLI2 and MLI1 firing rates as a function of distance from the activated CF. **j.** Average MLI2 and MLI1 responses evoked by (*left*) airpuff-evoked activation of CFs and MFs, (*middle*) activation of CF alone, and (*right*) activation of MF alone. **k.** Summary of firing-rate changes of individual MLIs. **l.** Simulations of airpuff evoked MLI1 responses that showed the largest increases and decreases in firing for simulations. **m.** As in **i** but for experimental responses. **n.** MLI1 responses (aligned to a PC_CS_) evoked by a single CF and by pairs of nearby CFs within 5 ms of each other *in vivo*. **o.** As in **n** but for simulations. **p.** Simulations of airpuff-evoked responses of MLI1s evoked by a single CF, and by 2, 5, and 10 nearby CFs. **q.** As in **o** but for individual MLI1s. **r.** (*left*) Schematic showing strong CF-MLI2 and weak CF-MLI1 connections. (*right*) Summary of the effects of spontaneous CF activation, airpuff-evoked responses for a single CF and for synchronous CF activation.

To gain insight into how the CF and MF pathways interact at the level of MLIs, we performed simulations based on large scale EM reconstructions of the molecular layer of the cerebellum^8,34^ (**Fig. 4f**), and included MLI2s, electrically-coupled MLI1s, PCs and CFs. The firing properties of individual cells, the synaptic properties, the electrical coupling between MLI1s and CF activation of MLIs were constrained by a combination of slice electrophysiology experiments, *in vivo* electrophysiology studies and EM reconstructions^8,15,34^ (**Extended Data Table 1**). Spontaneous CF activation increased the firing of MLI2s and inhibited MLI1s (**Fig. 4g**) as observed experimentally (**Fig. 3fg**). Simulations also allowed us to assess the spatial pattern of MLI2 excitation and MLI1 inhibition. The color-coded responses of MLI2s and MLI1s shows that a CF strongly excites several MLI2s in a parasagittal plane, and somewhat more broadly inhibits MLI1s (**Fig. 4h**), and this is also apparent in a plot of the average MLI1 and MLI2 firing frequency as a function of distance from the plane containing the activated CF (**Fig. 4i**). These simulations recapitulate how CFs influence MLI activity within a restricted parasagittal plane to regulate local circuitry within the molecular layer.

We next simulated airpuff-induced responses by activating CFs and granule cells with time courses guided by our in vivo recordings. Airpuffs evoke strong MLI2 excitation and weak average changes in MLI1 firing rates (**Fig. 4j**, *left*) that are similar to the firing rate changes observed *in vivo* (**Fig. 4de**, *left*). We evaluated the roles of MFs and CFs in these responses, and find that CF activation alone strongly excites MLI2s and inhibits MLI1s (**Fig. 4j**, *middle*), and that MFs excite both MLI2s and MLI1s (**Fig. 4j**, *right*).

The distribution of responses of individual cells shows that CFs alone inhibit most MLI1s, MFs alone excite most MLI1s, and coactivation of MFs and CFs, as with an airpuff, excites about half of the MLI1s and inhibits the others (**Fig. 4k**). For simulated responses of MLI1s evoked by airpuffs (CF + MF), a comparison of the MLI1 responses for the cells with the top and bottom 10% firing rate changes shows the wide range of MLI1 responses (**Fig. 4l**). Qualitatively similar results were observed for airpuff-evoked responses *in vivo* (**Fig. 4m**). Thus, we find that airpuffs increase the firing rates of some MLI1s primarily as a result of MF-Grc-MLI1 excitation, and decrease the rates of other MLI1s primarily as a result of CF-MLI2-MLI1 inhibition. Because GABAergic inhibition locally regulates the amplitudes of CF-evoked calcium signals in PC dendritic branches^35^, this spatial gating suggests that if the CF-MLI2-MLI1-PC pathway dominates for a given dendritic region it will promote plasticity of local GrC-PC synapses, but if the MF-MLI1-PC pathway dominates then plasticity of GrC-PC synapses will be suppressed.

## Synchrony of CF inputs biases towards disinhibitory pathway

What conditions might favor the CF-MLI2-MLI1-PC disinhibitory pathway to promote plasticity? Previous studies have shown that synchronous CF activation occurs *in vivo* in parasagittal bands under a variety of experimental conditions and is increased for sensory-evoked responses^36–38^. Synchronous optogenetic activation of CFs is highly effective at inducing cerebellar dependent learning^3^. We therefore explored how synchronous CF activation of nearby PCs influences MLI1 firing. We find that, on occasion, airpuffs elicit synchronous PC_CS_ (within 5 ms) that led to a stronger suppression of nearby MLI1s than when a PC_CS_ happened in isolation (**Fig. 4n**), and simulations replicated the stronger suppression of MLI1 firing for synchronous CF activation (within 5 ms) (**Fig. 4o**). It was challenging to record from neighboring PCs that received airpuff input for extended periods, so we also used simulations to assess the influence of CF synchrony on airpuff-evoked responses. We find that synchronous CF activation of nearby PCs leads to much stronger suppression of MLI1 firing, and the extent of the suppression increases as the number of synchronously firing PCs increases (**Fig. 4p**). Similarly, the fraction of MLI1s strongly suppressed also increases with the number of synchronously firing PCs (**Fig. 4q**). This suggests that synchronous activation of nearby CFs is highly effective at suppressing MLI1 firing, consistent with promoting plasticity (**Fig. 4r**).

These results predict that synchronous CF activation will disinhibit nearby PCs and increase the amplitude of CF-evoked dendritic calcium signals. We therefore used multiphoton imaging to directly measure CF-evoked calcium signals in PCs^36,39^(**Fig. 5a,b**) and compared their amplitudes for responses evoked by an isolated CF or when nearby CFs were synchronously activated. Spontaneous CF-evoked calcium increases were observed at 1-2 events/s (**Fig. 5c**, *top trace first event*). Sensory stimulation fails to evoked calcium increases in 63% of trials, evokes single events in 29% of trails, and evokes multiple events in 8% of trials (**Fig. 5c, e, f**), consistent with previous work showing that sensory stimulation can result in multiple short-latency CF spikes, large dendritic calcium signals and promote long-term synaptic depression at PF-PC synapses^32^. We compared CF-evoked calcium signals for spontaneous events, airpuff trials that evoked a single event, and airpuff trials that evoked multiple events (**Fig. 5e,g**). For trials in which the airpuff evoked multiple CF spikes the calcium signal is prolonged (**Fig. 5e**) and peak calcium signals are much larger (**Fig. 5e,g**). Consistent with previous reports, average calcium signals were also larger for airpuff trials that evoked a single CF spike than for spontaneous inputs ^31,32^ (**Fig. 5e, g**).

**Figure 5.**
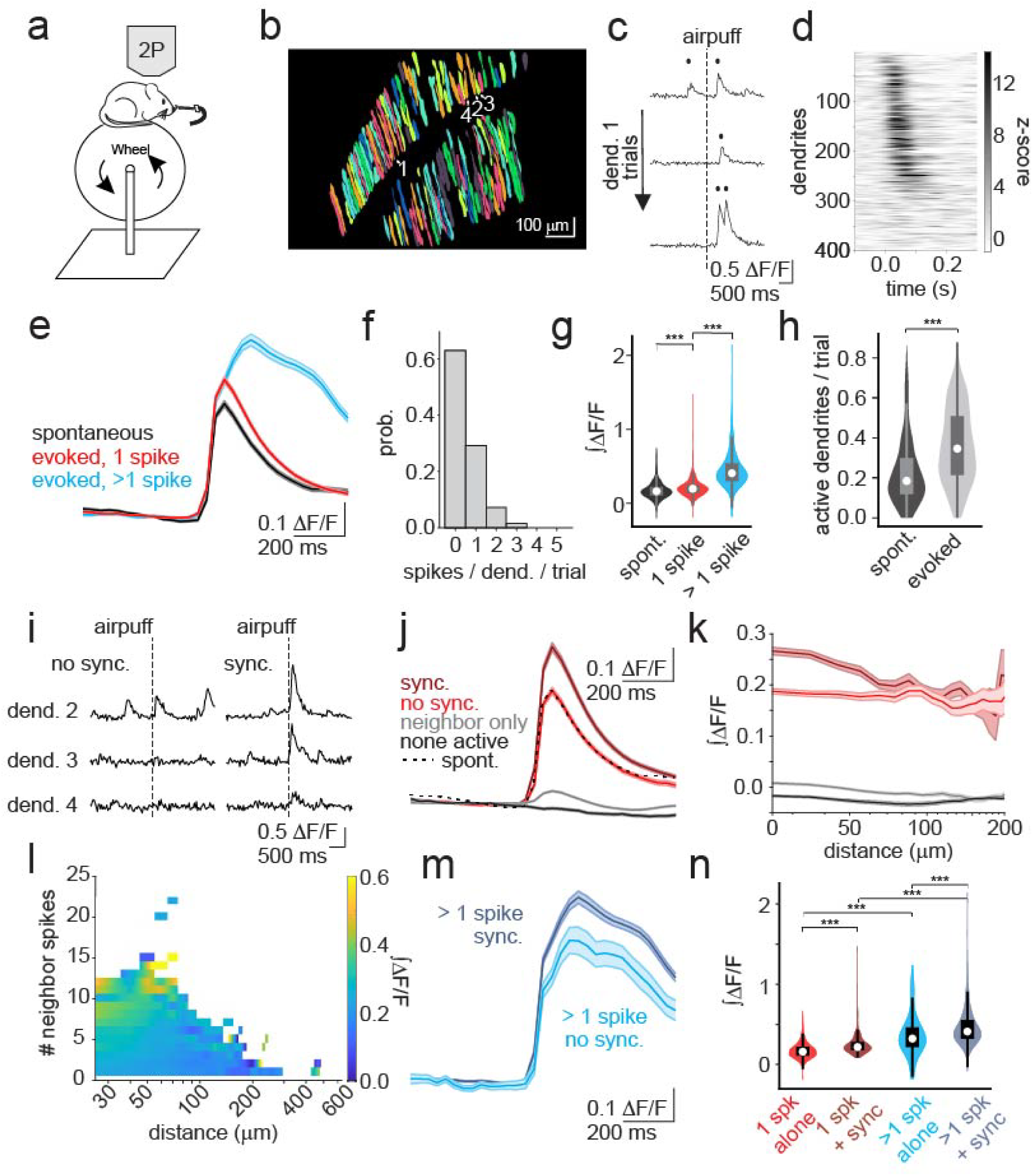
Sensory-driven CF synchrony enhances PC dendritic calcium signaling. **a.** Experimental design. Mice were placed on a freely moving wheel and 2P imaging of CF input to Purkinje cell dendrites was performed during ipsilateral airpuff delivery. **b.** Representative field of view of the extracted Purkinje cell dendritic regions of interest. Numbered dendrites correspond to the example data in (**c**,**i)**. **c.** Airpuff evoked calcium transients for an example dendrite in trials with one evoked deconvolved spike (PC_CS_) (top, middle) and two evoked deconvolved spikes (bottom). Filled circles represent spike times. **d.** Z-scored heatmap of airpuff evoked spikes sorted by latency to peak for each dendrite. **e.** Trial averaged calcium transients aligned to spontaneous spikes prior to an airpuff (black), single airpuff evoked spikes (red) and multiple airpuff evoked spikes (blue). **f.** Probability distribution of the number of airpuff evoked spikes that a dendrite fires per trial. **g.** Distributions of the integrated calcium transients of the conditions in (**e**) (spont. vs 1 spike, p=2.4e-06; 1 spike vs >1 spikes, p=3.6e-91). **h.** Distributions depicting the fraction of dendrites that fire at least one spike in a 500 ms window before the airpuff (spont.) and 500 ms after the airpuff (evoked) (p=4.2e-32). **i.** Example trials of airpuff evoked calcium transients of a reference dendrite and 2 neighbor dendrites that fire synchronously (right) or show no synchrony (left). Dendrites are highlighted in (**b**). **j.** Single spike trial-averaged calcium transients aligned to the spike of the reference dendrite when it fires in conjunction with neighbor dendrites (sync, dark red) up to 80 µm away, alone (no sync., red), or when only a neighboring dendrite is active (neighbor only, grey). When no dendrites were active (none active, black), the calcium transients were aligned to airpuff onset. Dotted line represents the mean spontaneous spike from (e). **k.** Integrated calcium transients of the reference dendrite for the conditions in (**j**), plotted as a function of distance between the reference dendrite and the neighbors. **l.** Heatmap depicting the calcium transient amplitude as a function of distance between the reference and neighbor dendrites and the number of spikes fired by the neighbor dendrites for the sync condition. **m.** Same as (**j**) but for trials in which the reference dendrite fire more than one spike. **n.** Distributions of the integrated calcium transients of the conditions in (**j**) and (**m**) (1 spk alone vs 1 spk + sync, p=8.3e-14; >1 spk alone vs >1 spk + sync, p=1.8e-05; 1 spk alone vs >1 spk alone, p=6.9e-20; 1 spk + sync vs >1 spk + sync, p=1.5e-52).

Airpuffs evoked responses in a large fraction of PC dendrites (**Fig. 5dhi**), which allowed us to assess the effect of synchronous CF activation (at least one response in nearby dendrites within ± 33 ms) on calcium signals (**Fig. 5j-l**). For airpuff-evoked responses comprised of a single event (**Fig. 5j**), synchronous activation of nearby CFs is associated with larger calcium transients compared to those evoked by either well-isolated CF activation or spontaneous events (**Fig. 5j**). It is unlikely that this enhancement reflects fluorescence contamination from neighboring dendrites, as it is also observed when fluorescence measurements are restricted to the brightest regions in the middle of each dendrite (**Extended Data Fig. 4**). We also observed a small elevation of calcium when only neighboring dendrites are active, consistent with disinhibition of the primary PC dendrite (**Fig. 5j**). We did not, however, observe any response when neighboring CFs are inactive (**Fig. 5j**), consistent with the observation that optogenetic silencing of CFs abolishes calcium signals in PC dendrites^32^ and the conclusion that GrC input does not produce measurable PC dendritic calcium transients. Notably, calcium signals evoked by well-isolated sensory CF activation and spontaneous CF activation are virtually identical (**Fig. 5j**), suggesting that the synchrony of neighboring CFs is responsible for the enhancement of average sensory-evoked responses. As predicted by our simulations of disinhibition, the influence of CF synchrony is minimal at distances above approximately 60 μm (**Fig. 5k**), and enhancement of CF-driven calcium transients is more pronounced when many nearby CFs are activated (**Fig. 5l**). We also find that synchronous activation of nearby CFs enhances calcium signals on trials where airpuffs evoke more than one event in the dendrite of interest (**Fig. 5m,n**). These results indicate that synchronously active CFs can effectively recruit disinhibition to significantly enhance PC dendritic calcium.

We conclude that CFs preferentially activate MLI2s via spillover, thereby leading to inhibition of MLI1s and disinhibition of PCs. We have shown through modeling and *in vivo* recordings that this newly identified CF-MLI2-MLI1-PC circuit is suited to enhance CF signals in PC dendrites, and that synchronous CF activation is particularly effective at engaging this circuit to elevate dendritic calcium signals in a manner suited to promote plasticity. This provides a powerful level of regulation that can account for the observation that disinhibitory mechanisms are important for cerebellar plasticity^20^, and that can explain why sensory stimuli or optogenetic manipulations that lead to widespread, synchronous CF activation^31,3^ are highly effective at producing cerebellar dependent plasticity and learning. Overall, this circuit motif allows CFs to act in a dual role by simultaneously providing direct PC excitation to trigger regenerative dendritic calcium spikes while also temporarily relieving synaptic inhibition to enhance the calcium transient associated with these spikes, thereby creating a brief temporal window for the enhanced calcium signaling known to be essential for inducing LTD.

## Methods

### Serial EM reconstructions

We previously imaged and aligned a 770 μm X 750 μm X 53 μm volume of lobule V of the mouse cerebellum for EM reconstructions comprised of 1176 45-nm thick parasagittal sections. We used automated image segmentation to generate neuron boundaries^34^. We used the neuron segmentation to reconstruct 10 CFs which were identified based on their characteristic morphology such as their large axons projecting into the molecular layer and running along individual PC. CFs were excluded if they were unable to be traced back to the molecular layer or if their respective PC was cut off.

CF spillover contacts onto MLIs were identified by the physical contact of the CF bouton to an MLI dendrite with vesicles present (typically from a CF-PC synapse) but a lack of clustering of vesicles toward the MLI membrane and the absence of a post synaptic density. MLI contacts were analyzed for 10 CFs and subtyped into MLI1 or MLI2 based off of previous identification methods^8^. The surface areas of CF contacts onto MLIs were determined by tracing the contact area using annotation tools in Neuroglancer, retrieving their 3D coordinates and through MATLAB calculating the 3D surface area. Parallel fiber synapses onto MLIs were determined by their small diameter boutons and completely parallel morphology which was traced through the depth of the dataset.

CF-MLI1-PC and CF-MLI2-MLI1-PC synapses onto local and neighboring PCs were identified using automated synapse prediction. The neighboring PC was determined to be the PC immediately adjacent to a PC that has an identified CF. A total of 608 CF-MLI1-PC synapses were made onto local PCs and 388 onto neighboring PCs while a total of 2799 CF-MLI2-MLI1-PC synapses were made onto local PCs and 2583 of those synapses onto neighboring PC. Pathway analysis was done using python.

### Slice experiments

Animal procedures were performed in accordance with the NIH and Animal Care and Use Committee (IACUC) guidelines and protocols approved by the Harvard Medical School Standing Committee on Animals. C57BL/6 mice were obtained from Charles River Laboratories. Animals of either sex were randomly selected for experiments. Animals were housed on a normal light–dark cycle with an ambient temperature of 18–23 °C with 40–60% humidity.

Acute parasagittal slices (220-μm thick) were prepared from P28-45 C57BL/6 mice. Mice were anaesthetized with an intraperitoneal injection of ketamine (10 mg kg^−1^) and perfused transcardially with an ice-cold solution containing (in mM): 110 choline chloride, 7 MgCl_2_, 2.5 KCl, 1.25 NaH_2_PO_4_, 0.5 CaCl_2_, 25 glucose, 11.6 sodium ascorbate, 3.1 sodium pyruvate, 25 NaHCO_3_, equilibrated with 95% O_2_ and 5% CO_2_. Slices were cut in the same solution, and then transferred to artificial cerebrospinal fluid (ACSF) containing (in mM) 125 NaCl, 26 NaHCO_3_, 1.25 NaH_2_PO_4_, 2.5 KCl, 1 MgCl_2_, 1.5 CaCl_2_ and 25 glucose equilibrated with 95% O_2_ and 5% CO_2_. As indicated, some experiments were performed in elevated external calcium (2.5 mM). Following incubation at 34 °C for 30 min, the slices were kept up to 6 h at room temperature until recording.

Recordings were performed at 32 °C with an internal solution containing (in mM): 35 CsF, 110 CsCl, 10 HEPES, 10 EGTA and 2 QX-314 (pH adjusted to 7.2 with CsOH, osmolarity adjusted to 300 mOsm kg^−1^). Visually guided whole-cell recordings were obtained with patch pipettes of ∼3-MΩ resistance pulled from borosilicate capillary glass (BF150-86-10, Sutter Instrument). Electrophysiology data were acquired using a Multiclamp 700B amplifier (Axon Instruments), digitized at 20 kHz and filtered at 4 kHz using Igor Pro (Wavemetrics) running mafPC (courtesy of M.A. Xu-Friedman). Acquisition and analysis of slice electrophysiological data were performed using custom routines written in Igor Pro (Wavemetrics). The following receptor antagonists were added to the ACSF solution to block GABAergic and glycinergic synaptic currents (in μM): 10 SR95531 (gabazine), 1.5 CGP, and 1 strychnine. All drugs were purchased from Abcam and Tocris.

We recorded MLIs in voltage clamp at −65mV. Recordings were made from lobules IV-V of the vermis. We recorded from MLIs in the inner two-thirds of the molecular layer and determined the identity of MLI1s and MLI2s by characterizing their characteristic electrical properties as previously described^8^. We used spikelets to identify MLI1s, and a lack of spikelets combined with a high input resistance to identify MLI2s.

We used a theta glass stimulus electrode to stimulate CFs in the granule cell layer with pairs of stimuli (350 µs in duration, 50-ms ISI, 10-100 μA, 10-s intertrial interval) at different locations in an area ∼50 µm into the granule cell layer and ∼100 µm on each side of the soma of the MLI being recorded. We recorded >10 sites per cell in search of inputs that were all-or-none and with marked paired-pulse depression (PPR<0.6). We then reduced the stimulus intensity to threshold to evaluate the all-or-none nature of the response and contamination from granule cell inputs. Stimulation at the threshold for CF activation stochastically evokes successes and failures, and slight increases in stimulus intensity eliminate failures. This indicates it is a single all-or-none input. To evaluate if the CF input was isolated, failure trials were examined for granule cell responses. Traces shown are averages of 10 trials. For analysis of the kinetics, we calculated the 20-80 rise and decay times. We bath applied dl-threo-β-benzyloxyaspartic acid (DL-TBOA; 50 μM) while stimulating CF inputs with single stimuli every 10 s. The access resistance and leak current of the MLIs were monitored continuously. Traces shown are averages of 10 trials for baseline and in the presence of TBOA. Parallel fibers were stimulated in the molecular layer with pairs of stimuli (350 µs in duration, 140–250 μA, 20-ms ISI, 5-s intertrial interval) within ∼100 µm of the soma of the MLI being recorded. Parallel fiber inputs with paired-pulse facilitation were found for all MLIs recorded.

We characterized the contributions of AMPARs and NMDARs to CF-MLI and PF-MLI synaptic currents with pharmacology. We recorded EPSCs from MLIs at +40mV with a 10-s intertrial interval. Voltage steps were 3.5 s in duration, and single stimuli were delivered at 2.5 s. Voltage steps without stimulation were recorded each trial for subtraction. We then blocked the AMPA component with NBQX (5 µM) and followed this by bath application of the NMDAR antagonist CPP (2 µM). The access resistance and leak current of the MLIs were monitored continuously. Current traces are the average of approximately 10 trials. The NMDA/AMPA ratio was calculated as the peak current of the NMDA component divided by the peak current of the AMPA component, with both components measured at +40 mV.

### HCR-FISH

Acute cerebellar slices from p28-p45 mice were prepared as described, and fixed for 2 hours in 4% paraformaldehyde in PBS (Biotum) at 4 °C. Slices were stored overnight in 70% ethanol in RNase-free water at 4 °C. A floating slice HCR protocol^15^ was performed with the following probes and matching hairpins (Molecular Instruments): sortilin related VPS10 domain containing receptor 3 (*Sorcs3*), neurexophilin 1 (*Nxph1*), and glutamate receptor, ionotropic, NMDA2B (epsilon 2) (*Grin2b*). MLI1s express *Sorcs3*, and MLI2s express *Nxph1*. Amplification hairpins were B1-750, B2-488, and B3-647. Slices were mounted on slides (Superfrost Plus, VWR) with mounting medium (Fluoromount, ThermoFisher) and no.1 coverslips. Images were acquired with a Leica Stellaris X5 confocal microscope using a 63x oil immersion objective (1.4 NA, Olympus). The reporter and HCR probe/hairpin channels were imaged with 180 nm resolution in a 20-µm thick, 1-µm interval tiled z series in lobule IV/V. *Sorcs3*+ and *Nxph1*+ cell locations in the molecular layer were manually labelled using the multi-point tool in Fiji. *Grin2b* fluorescence in each cell was averaged within a 7-µm diameter circular mask in Matlab (Mathworks).

### In vivo electrophysiology and calcium imaging

Mice for *in vivo* electrophysiology recordings (wild type C57BL/6J) and calcium imaging (Tg(PCP2-Cre)3555Jdhu) were used in accordance with approval from the Duke University Animal Care and Use Committee. Animals were housed on a normal light–dark cycle, and animals of either sex were randomly selected for experiments.

Mice were anesthetized and placed on a stereotaxic apparatus. Isoflurane was delivered at 5% for induction and at 1.5% for maintenance throughout surgery. A titanium headpost (HE Palmer) was secured to the skull with metabond (Parkell). Body temperature was maintained with a heating pad (TC-111 CWE). Bupranex (0.05 mg/kg, subq) was administered for 24 h after surgery and animals were monitored for 4 days.

For in vivo electrophysiology experiments, after 2+ weeks of recovery, mice were anesthetized with 2% isoflurane and a small craniotomy (1 mm) was drilled over the primary fissure of lobule simplex. The craniotomy was covered with Qwik-Cast (WPI) and a thin layer of metabond between recording days. 1-week post-surgery, mice were head fixed and placed on a motorized circular treadmill for 45 minutes over 4 days. The rotation speed was initially set at 0.06 m/s and was increased in steps of 0.02 m/s per day until reaching a final speed of 0.12 m/s. After this initial phase, 2 more days were added in which the motor was turned off and on in alternating ∼10 second epochs while mice were on the treadmill. Following habituation, the craniotomy was opened and a Neuropixels 1.0 probe coated with dye (DiI or DiD) was lowered at a rate of 3 μm/sec into the right primary fissure of the cerebellum (AP: −5.82, ML: 2.0 from bregma) reaching a final placement of ∼2000 μm below the cortical surface. The probe was allowed to settle for 45 minutes before data collection. A blunted 23-gauge needle (PrecisionGlide) was placed 3 mm from the right eye of the mouse to deliver corneal airpuffs (25 psi). Air flow was controlled with a solenoid (The Lee Company) and an air pressure regulator (SNS). 100 airpuffs were delivered during running or stationary epochs, and 100 airpuffs were delivered during motor on or running epochs. Mice were recorded for 2 or 3 days (1 session each day. 3 mice, 6 sessions). After the last day of recording, mice were euthanized and perfused for histology. 50 μm brain sections were collected to visualize and confirm the probe location. Electrophysiology data were collected with SpikeGLX.

For imaging experiments, a craniotomy (3 mm) was drilled over lobule simplex of the right cerebellar hemisphere. For GCaMP expression targeted at PC dendrites, a glass micropipette was filled with virus AAV1.CAG.Flex.GCaMP6f.WPRE.SV40 (UPenn vector core, titer = 1.2 × 10^13^) or AAV1.CAG.Flex.GCaMP7f.WPRE.SV40 (Addgene 104496, titer = 1.5 × 10^13^) diluted 1:4 – 1:20 in ACSF (titer for injection: 7.5 × 10^11^). 150-200 nL of virus were injected at 2-4 sites in the craniotomy. The injection rate was 30-50 nL/min.

Two-photon imaging was performed using a resonant scanning microscope (Neurolabware) with a 16x water immersion objective (Nikon CFI75 LWD 16xW 0.80NA). To stabilize the immersion solution for imaging, a polymer (MakingCosmetics, 0.4% Carbomer 940) was used. For GCaMP imaging, A 920 nm wavelength Ti:Sapphire laser was raster scanned via a resonant galvanometer (8 kHz; Cambridge Technology) onto the cerebellum at a frame rate of 30 Hz with a laser power of <40 mW and a field of view of 1030 μm x 581 μm (796 x 264 pixels). Emitted photons were collected through a green filter (510 ± 42 nm band filter (Semrock)) onto a GaAsP photomultiplier (H10770B-40, Hamamatsu) and the microscope was controlled using Scanbox software (Neurolabware).

### Analysis and statistics

For in vivo Neuropixels recordings, units were sorted using Kilosort 2.0 and manually curated with Phy as described previously^8^. After unit curation, layer and cell type was identified using a deep-learning classifier^40^. Units were separated into PC simple spikes (PC_SS_), PC complex spikes (PC_CS_), mossy fibers, molecular layer interneurons (MLIs) and unclassified units. To obtain the initial population of putative MLIs, we pooled together units in the molecular layer that the classifier label as MLIs or unclassified units. After this initial step, manual MLI1 and MLI2 classification was performed as previously described in detail^8^.

For spontaneous CS CCGs, the peak changes in spike rates following the complex spike were measured as the averages of t_1_=1-2 ms and t_2_=3-6 ms for MLIs, and t_1_=0-1 ms and t_2_=4-7 ms for PCs. The cumulative changes in spike rates were measured as the average of 20-30 ms following the complex spike of the integrals of the CCGs. All responses were measured relative to baseline averaged 30 ms prior to the complex spike. Pairs were determined to be connected if the response z-score was >4 or <-4. Latencies were measured as the half-max times for t_1_ and t_2_ for connected pairs with response z-scores >10 or <-10.

Peristimulus time histograms were constructing by convolving the trial averaged activity of each unit with a 6ms gaussian kernel and then averaging across units. Traces then were aligned to airpuff onset. Mossy fiber and PCss heatmaps were cross validated using half of the trials to calculate the max airpuff response for each unit and then used to sort the averaged z-scored responses of the remaining trials. MLI1 and MLI2 response heatmaps were constructed by averaging the z-scored activity of all MLIs connected to the units in the PCcs heatmap. MLIs without a connected PCcs are blanked (white lines). Significantly airpuff driven cells were obtained by comparing the activity 100ms before and after the airpuff using the Wilcoxon signed rank test. Data are reported as mean ± standard error, and statistical significance was defined as P<0.05. * = P<0.05. ** = P<0.01. *** = P<0.001.

For in vivo calcium imaging analysis, raw data was motion corrected on the X and Y planes using sub-pixel image registration. PC dendrite isolation was performed with a post-hoc processing pipeline consisting of principal component analysis (PCA) followed by individual component analysis (ICA) (Mukamel et al. 2009). Final dendrite segmentation was achieved by applying a gaussian filter and thresholding pixels of individual components. A binary mask was created by combining highly correlated pixels (correlation coefficient > 0.80) and removing overlapping regions between segmented dendrites. Any PC somas present in the field of view were avoided. Raw calcium traces were filtered (high pass Butterworth filter, 0.035 Hz, order 3) and the MATLAB OASIS package^41^ was used to identify events corresponding to complex spikes. The ‘FOOPSI’ method was used to account for the shape of the Ca^2+^ transients using GCaMP as a Ca^2+^ indicator. In addition, a threshold of 3 standard deviations above baseline was used to identify transients.

ΔF/F signals were measured on a trial-by-trial basis by using the mean of a 500 ms window prior to each airpuff as F0. Spikes that occurred in a 500ms window 2 seconds before the airpuff were classified as spontaneous spikes. Spikes that occurred between 0 and 500 ms after airpuff onset were classified as airpuff evoked spikes. In both conditions, when there was an additional spike within 500ms of the first spike, the instance was classified as two (or more) spikes. Active dendrites were defined as dendrites that fire at least one spike in the ‘spontaneous’ window or the airpuff window and their distributions were compared using the Mann-Whitney U test. PSTHs were constructed by aligning DF/F traces to the first spike after airpuff onset and averaging them across trials. For spontaneous spikes, DF/F traces were aligned to the first spike in the equivalent 500 ms window 2 seconds prior to airpuff onset. Spiking heatmaps were constructed using cross validated deconvolved spike PSTHs aligned to airpuff onset. Cross validation was performed using half of the trials to calculate the latency to the max airpuff response for each dendrite and then using that order to sort the averaged z-scored responses of the remaining trials. For spike synchrony analysis, synchrony was defined as the co-firing of two or more dendrites within a temporal window of 66 ms (one frame before or after the reference spike). To assess the effect of distance between synchronous dendrites on calcium influx, a sliding window of 50 µm with 10 µm steps was used. Trials were classified as synchronous when two or more dendrites co-fired in the time and distance windows, and they were classified as not synchronous otherwise. Trials in which two or more dendrites closer than the target distance window co-fired were removed from analysis. For centroid ROI analyses, the centroid was defined as the pixel located in the median x, y coordinate of the dendrite and one adjacent pixel in each direction, making a total of 9 pixels (∼4 µm²). Therefore, centroid fluorescence was the averaged value of those 9 pixels. For assessing statistical significance, DF/F traces in the 500ms window were integrated for each condition and compared using the Kruskal-Wallis with Dunn’s test post-hoc. Data are reported as mean ± standard error and statistical significance was defined as P<0.05. * is P<0.05. ** is P<0.01. *** is P<0.001.

### Simulations

The model of the molecular layer of the cerebellar cortex consisted of 840 MLI1s, 315 MLI2s, and 170 PCs with their associated CF inputs (**Fig.4f**). The model was constrained by EM reconstructions, slice recordings, and *in vivo* recordings of firing properties. The distributions of MLI1, MLI2 and PC baseline firing rates, capacitances and input resistances were based on slice and *in vivo* studies^8,15^. Synaptic connections made by MLI1s and MLI2s were based on slice recordings and EM reconstructions^8^. MLI1s were electrically coupled to each other with properties based on slice recordings. Model neurons were placed in the volume, and dendritic and axonal arbors assigned as three-dimensional right rectangular prisms centered at the somas, with dimensions based on EM reconstructions. Synaptic and electrical connections were then made between MLIs with probabilities and strengths based on the volume of arbor overlap (axodendritic for synaptic connections, dendrodendritic for MLI1-MLI1 electrical connections), scaled by measurements of electrical coupling in slice experiments. CF-MLI1 and CF-MLI2 synaptic connections were based on a combination of EM reconstructions and slice experiments. Inputs due to MF activity were modeled as granule cell inputs to MLIs with spike times drawn as Poisson events and their kinetics based on slice recordings.

MLIs were modeled as single-compartment exponential integrate-and-fire (EIF) neurons with membrane potential (*V*) dynamics obeying

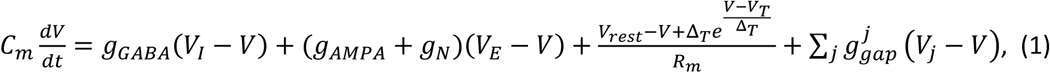

where *C*_*m*_ and *R*_*m*_ are the membrane capacitance and resistance, *g*_*GABA*_ and *g_AMPA_* are the inhibitory and excitatory conductances with reversal potentials *V*_*I*_ and *V*_*E*_, *g*_*N*_ is the conductance noise, *V*_*rest*_ is the resting potential, *V*_*T*_ is the threshold, Δ_*T*_ is the spike slope factor, and 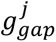 is the gap junction conductance with an electrically coupled cell *j* with membrane potential *V*_*j*_. Parameter values and definitions are given in **Extended Data Table 1**. Simulations used the Euler methods with 10 μs time steps. Synaptic currents were modelled as a difference of exponentials with rise times, decay times and latencies determined from experimental data. For MLI1s an electrical coupling term mediated by dendrodendritic gap junctions was added that was proportional to the somatic voltage difference between electrically coupled cells. An exponential spike-generating current was added^42^ to reproduce quantitatively MLI1-MLI1 crosscorrelations observed in slice experiments^8^. This current described the initiation and rising phase of action potentials, using two parameters, a soft threshold *V*_*T*_ and a spike slope factor Δ_*T*_. When the membrane potential reached its peak *V*_*θ*_, the dynamics in the equation 1 were replaced with a linear downward ramp towards the reset potential lasting 1 ms. Following each action potential, there was a refractory period drawn randomly from a uniform distribution, during which the membrane potential evolved freely under the dynamics in equation 1 with the spike-generating current disabled.

For each of the 10 reconstructed CFs (**Fig. 1**), two sets of aggregate data were compiled from the verified contact locations obtained from the serial EM reconstructions. (1) The numbers of contacts onto each postsynaptic MLI were recorded along with its cell type (MLI1 or MLI2). (2) The spatial bounding boxes of these contacts along the three directions were recorded for each CF (min and max coordinates along the three directions, corrected to have the PCL at y=0). Using this aggregate data, model CFs were chosen randomly from the 10 reconstructed CFs with replacement and placed at the location of each PC soma. To determine its connections with MLI1s and MLI2s, the empirical list of contact numbers with cells of that postsynaptic cell type were assigned in order of axodendritic overlap volume. To convert from contact numbers to synaptic conductances we assumed that the conductance was proportional to the number of contact sites. We estimate the conductance corresponding to an individual contact for by dividing the maximal observed conductance by the maximum number of contacts observed in reconstructions. This yielded a spillover quantal size of 1.2 nS for MLI2s and 0.375 nS for MLI1s.

The network simulator along with its analysis was written in Python using the package ANNarchy^43^ and run on a 16-core machine. Simulations were prepared and run in batches with the same initial state and different run conditions. For example, during the stimulation of the 10 individual CFs, a batch of identical networks was prepared, while in each network a different CF was stimulated along with other cases such as the co-activation of MFs. This allowed individual neurons to be compared across different runs in the batch. Simulations consisted of a 60s baseline, followed by 5000 300 ms long airpuff trials, for a total of 1560 s simulated time.

## Data Availability

Datasets supporting the findings of this study are available from the corresponding author upon request. Upon acceptance of the manuscript a minimum dataset will be provided through deposition in a public repository.

## Acknowledgments

We thank Lindsey Glickfeld, David Herzfeld and Gord Fishell for comments on the manuscript. WGR thanks Jacques Wadiche for helpful discussions. This work was supported by the NIH (R35NS097284 to W.G.R., F32NS133036 to E.P.L., R21NS085320, RF1MH128949, and RF1MH114047 to W.-C.A.L.,, R01NS128054 and R01NS112917 to C.A.H.), the Ruth K. Broad Biomedical Research Foundation (388000038 to M.E.H.), the Bertarelli Program in Translational Neuroscience and Neuroengineering, Stanley and Theodora Feldberg Fund, and the Edward R. and Anne G. Lefler Center. Portions of this research were conducted on the O2 High Performance Compute Cluster at Harvard Medical School.

---

**Extended Data Table 1.**
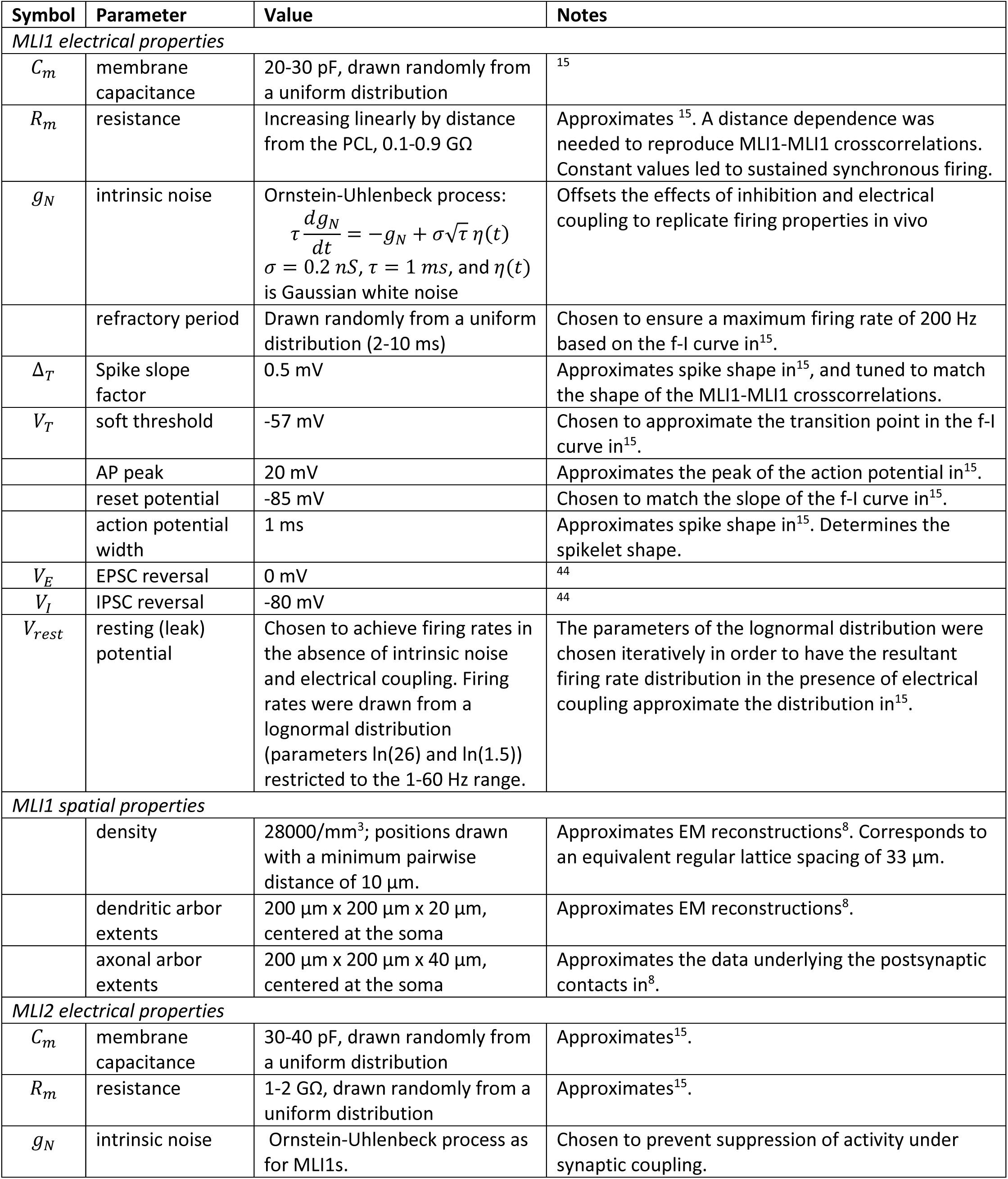
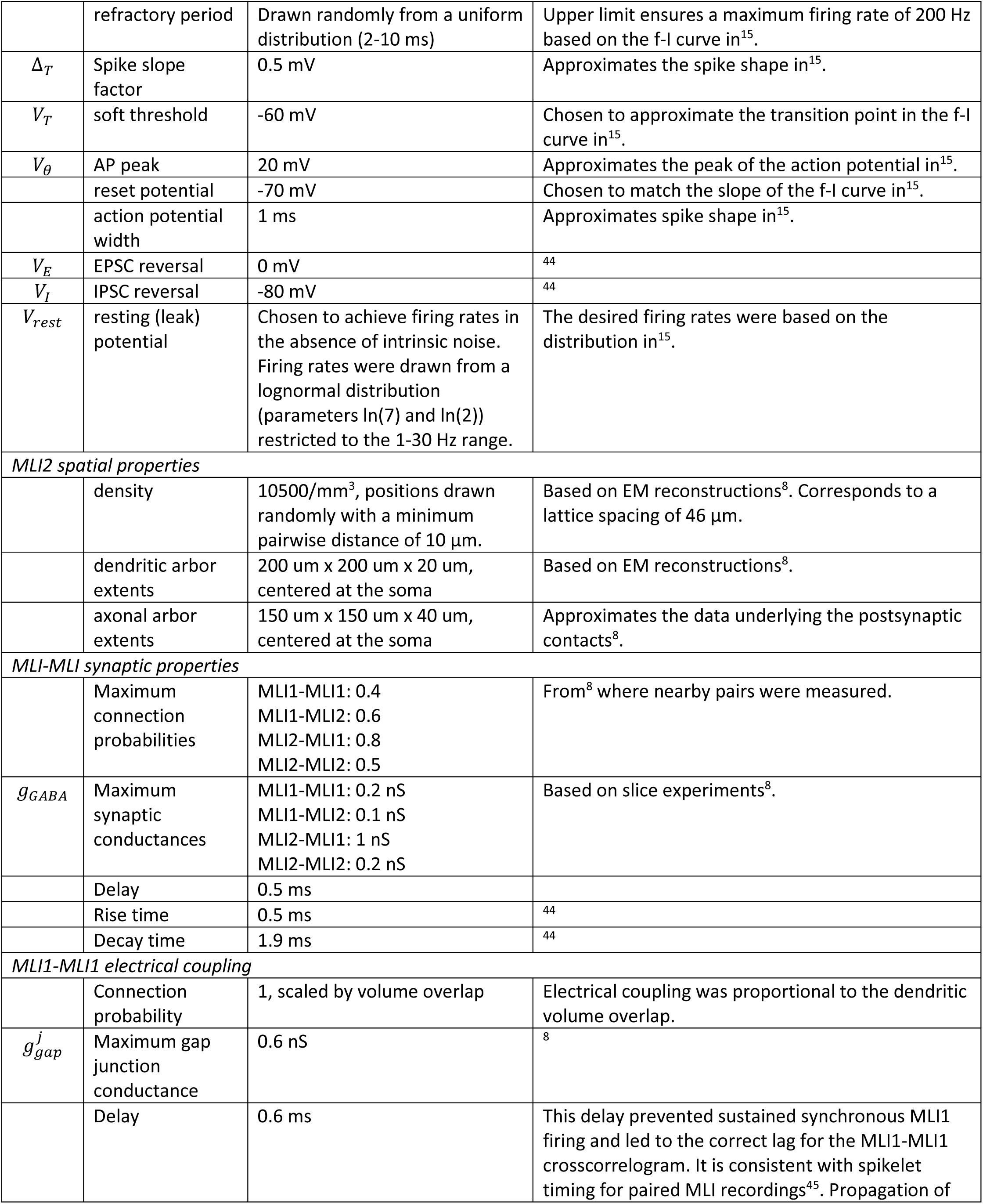
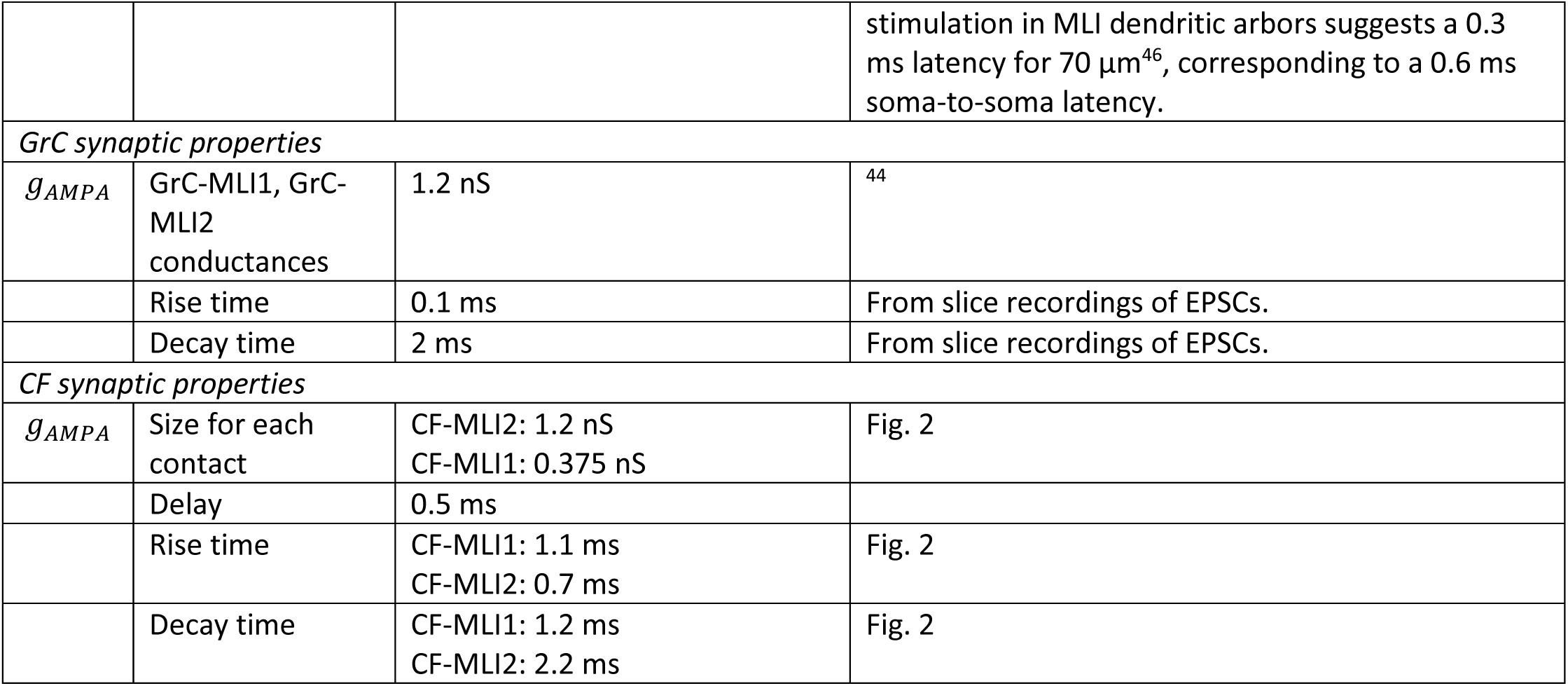
Model Parameters.

