## Supplemental Figures for "Climbing fibers selectively recruit disinhibitory interneurons to enhance dendritic calcium signaling in cerebellar Purkinje cells"

**Supplementary Figures**

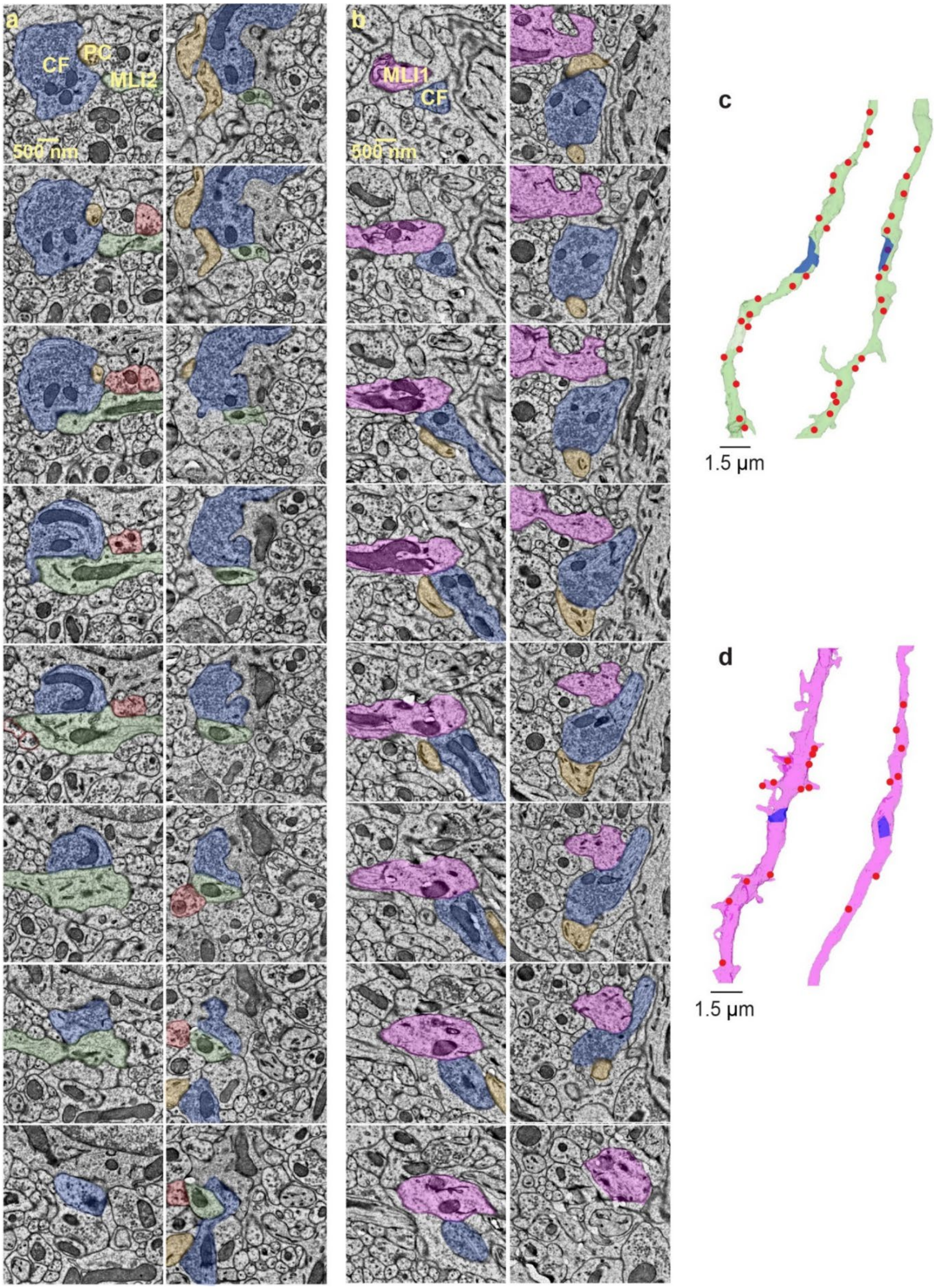

**Extended Data Figure 1|Examples of serial EM sections showing CF-MLI contacts.**

**a.** Series of EM sections shows a CF (*blue*) contacting an MLI2 (*green*) for two CF-MLI2 contacts (similar to **Fig. 2a**), with each column corresponding to a different CF-MLI2 contact.

**b.** As in a but for two CF-MLI1 contacts.

**c.** EM reconstructions based on images that include those in a. CF contact (*blue*) MLI2 (*green*) contact, and the red dots indicate the GrC-MLI contact sites.

**d.** As in c, but for CF-MLI1 contacts and the series in b.

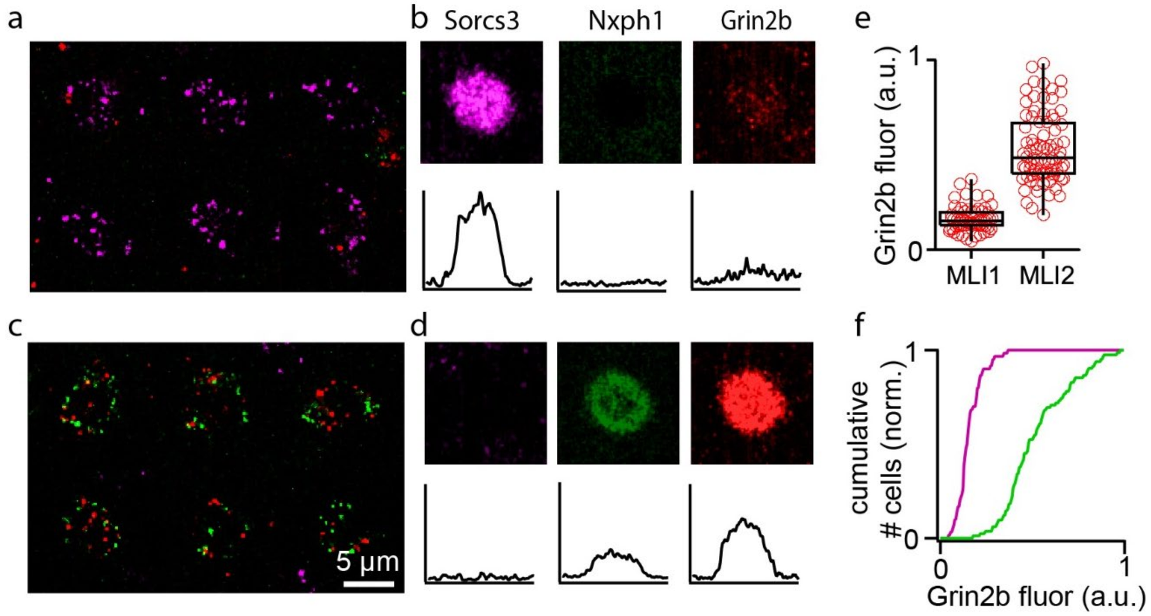


**Extended Data Figure 2|The NMDAR subunit *Grin2b* is present at much higher levels in MLIs than MLI2s.** Probes for *Sorcs3 (purple), Nxph1 (green) and Grin2b (red)* were labeled using HCR. *Sorcs3 (purple)* labels MLI1s, and *Nxph1 (green)* labels MLI2s.

**a.** MLI1s identified on the basis of prominent *Sorcs3* expression did not have prominent *Grin2b* expression.

**b.** Average fluorescence for *Sorcs3+* cells (n=50).

**c.** MLI2s identified on the basis of prominent *Nxph1 expression* also had clear *Grin2b* expression.

**d.** Average fluorescence for *Nxph1+* cells (n=50).

**e.** Summary of *Grin2b* fluorescence for individual MLI1s (*Sorcs3+*) and MLI2s (*Nxph1+*).

**f.** Normalized cumulative *Grin2b* fluorescence for individual MLI1s and MLI2s.

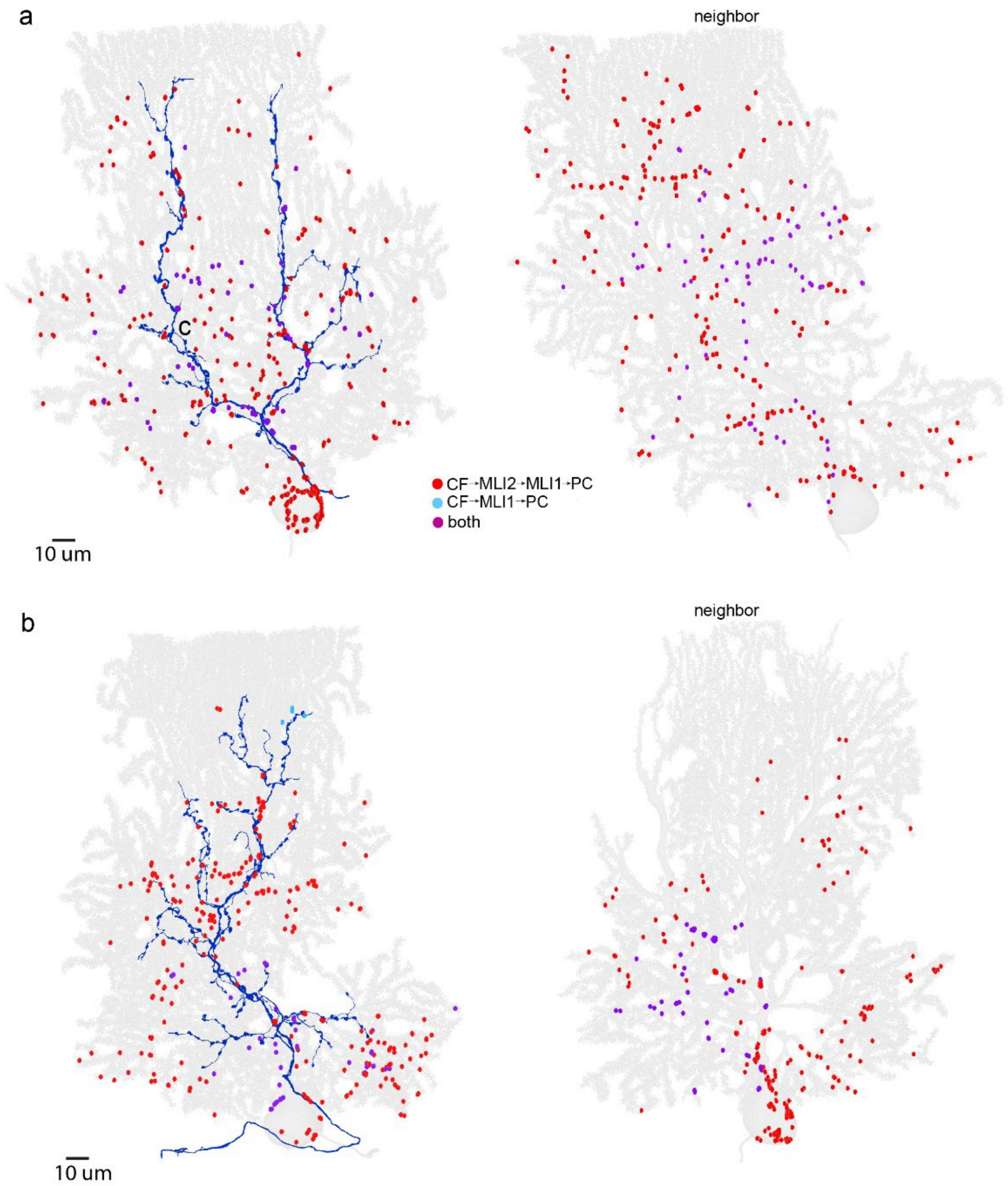


**Extended Data Figure 3|Examples of EM reconstructions used to evaluate the circuitry influenced by CF activation.**

**a.** Reconstructions are shown as in **Fig. 3c**. For a PC directly excited by a CF (dark blue), inhibitory synapses made by MLI1s that are contacted by the CF (CF-MLI1-PC1, light blue), MLI1-PC1 synapses that are CF-MLI2-MLI1-PC1 circuit (*red*), and MLI1-PC synapses for MLIs that are both directly contacted by CFs, and part of the CF-MLI2-MLI1-PC1 circuit (*purple*). (*right*) Reconstruction of a neighboring PC and the MLI contacts.

**b.** As in (**a**) but for a different pair of PCs.

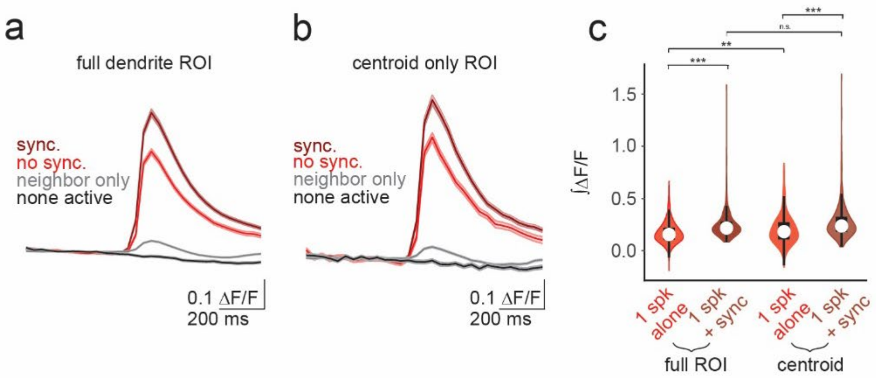


**Extended Data Figure 4| PC$_{CS}$ enhancement of calcium signaling during synchrony is not due to fluorescence bleed through at boundaries**

**a.** Trial averaged calcium transients from **Fig.5j**.

**b.** Same as A but averaging calcium transients at the dendrite's centroid. The centroid is defined as the pixel located in the median x,y coordinate of the dendrite and one adjacent pixel in each direction, making a total of 9 pixels (~4 μm²).

**c.** Distributions of the integrated calcium transients of the traces in A and B (Full ROI 1 spike alone vs Full ROI 1 spike + sync, p=6.8e-20; Centroid 1 spike alone vs Centroid 1 spike + sync, p=1e-14; Full ROI 1 spike alone vs Centroid 1 spike alone, p=5.6e-03; Full ROI 1 spike + sync vs Centroid 1 spike + sync, p=2.3e-01).
